# Recombinase-mediated integration of a multigene cassette in rice leads to stable expression and inheritance of the stacked locus

**DOI:** 10.1101/2020.04.16.045435

**Authors:** Bhuvan Pathak, Vibha Srivastava

## Abstract

Efficient methods for multigene transformation are important for developing novel crop varieties. Methods based on random integrations of multiple genes have been successfully used for metabolic engineering in plants. However, efficiency of co-integration and co-expression of the genes could present a bottleneck. Recombinase-mediated integration into the engineered target sites is arguably a more efficient method of targeted integration that leads to the generation of stable transgenic lines at a high rate. This method has the potential to streamline multigene transformation for metabolic engineering and trait stacking in plants. Therefore, empirical testing of transgene(s) stability from the multigene site-specific integration locus is needed. Here, the recombinase technology based on Cre-*lox* recombination was evaluated for developing multigenic lines harboring constitutively-expressed and inducible genes. Targeted integration of a 5 genes cassette in the rice genome generated a precise full-length integration of the cassette at a high rate, and the resulting multigenic lines expressed each gene reliably as defined by their promoter activity. The stable constitutive or inducible expression was faithfully transmitted to the progeny, indicating inheritance-stability of the multigene locus. Co-localization of two distinctly inducible genes by heat or cold with the strongly constitutive genes did not appear to interfere with each other’s expression pattern. In summary, high rate of co-integration and co-expression of the multigene cassette installed by the recombinase technology in rice shows that this approach is appropriate for multigene transformation and introduction of co-segregating traits.

**Significance Statement:** Recombinase-mediated site-specific integration approach was found to be highly efficacious in multigene transformation of rice showing proper regulation of each gene driven by constitutive or inducible promoter. This approach holds promise for streamlining gene stacking in crops and expressing complex multigenic traits.

## Introduction

The demand for resilient, productive, and value-added crops mandates breeding with multiple genes. Biotechnology innovates crop improvement through rapid introduction of genes; however, independently added genes are difficult to stack into cultivars. The population size needed to isolate a stacked F2 plant increases exponentially with the increasing number of unlinked genes, and screening of thousands of F2 plants becomes necessary for breeding 5 – 6 unlinked genes as compared to a handful of plants for a single gene. Therefore, breeding could be turned into a high efficiency process by simply linking the genes and stacking them into a locus (Petolino and Kumar, 2016; Que et al., 2010).

Multigene stacking could be obtained through co-bombardment of plasmids that tend to recombine and co-integrate into the plant genome; however, co-expression of the introduced genes, in this method, generally occurs at a low rate (Chen et al., 1998; Gelvin, 1998; Schmidt et al., 2008; Zhu et al., 2008). *Agrobacterium*-mediated T-DNA transfer is also an excellent method of multigene transformation as T-DNA harboring many genes enters into the plant cell and integrates into the genome (Collier et al., 2018; Li et al., 2003; Ruiz-Lopez et al., 2015). However, integration of tandem repeats or truncated copies of T-DNA, and disruption of critical genomic regions cannot be ruled out. These features of random transformation methods pose major challenges for multigene transformation, rendering a high number of the recovered events unsuitable for product development (Anand and Jones, 2018; Halpin, 2005).

The targeted integration approaches, on the other hand, are ideal for precise integrations into selected genomic sites. A number of targeted methods have been described that can be grouped into (a) recombinase-mediated integration into pre-characterized sites, and (b) integrations into double-stranded breaks (DSB) induced by site-specific DSB reagents such as CRISPR/Cas9 (Chen and Ow, 2017; Petolino and Kumar, 2016; Srivastava and Thomson, 2016; Weeks et al, 2016). Each method has its own advantages and disadvantages. While recombinase-mediated methods are highly efficient in targeted integrations, e.g., with Cre-*lox*, FLP-*FRT* and other site-specific recombination systems, they require placement of recombination target sites that can subsequently be targeted for gene stacking (Anand et al., 2019; Li et al., 2009; Ow, 2003; Srivastava and Gidoni, 2010). The CRISPR/Cas9 method, on the other hand, can target virtually any site in the genome, but the non-homologous mode of DSB repair in plant cells overrides the integration of exogenous DNA into the DSB site (Puchta and Fauser, 2014; Voytas, 2013). As a result, the current DSB-based approaches require extensive efforts and expanded post-transformation analysis for isolating the rare individuals that contain targeted integrations (Svitashev et al., 2015; Yang et al., 2019). Therefore, although, recombinase-mediated approach can only be practiced on the ‘prepared’ target lines, it provides a promising platform for developing gene stacking technologies for crops.

Cre-*lox* site-specific recombination is one of the most attractive systems for recombinase-mediated genome engineering. Its high efficiency has been demonstrated in a number of plant species, most of which develop into healthy fertile plants (Gidoni et al., 2008; Srivastava and Thomason, 2016). The phenotypic effects of Cre recombinase in plants are either undetectable or reversible owing to its high specificity and undetectable off-target activity (Coppoolse et al., 2003; Ream et al., 2005; Srivastava and Nicholson, 2006). Other site-specific recombination systems that show high efficiency in plant cells include FLP-*FRT*, phiC31-*att* and Bxb1-*att* systems (Anand et al., 2019; Hou et al, 2014; Li et al., 2009; Nandy et al., 2011; Ow, 2011; Thomson and Ow, 2006). Recombinase-mediated gene stacking has been demonstrated in soybean by inserting gene silencing and overexpression constructs through two rounds of transformation (Li et al., 2010). However, more research is needed to test the efficiency of gene stacking by recombinases, and verify its efficacy for co-expression of the stacked genes. Further, since the stacked genes could consist of conditionally or developmentally regulated genes, it is important to test the recombinase-mediated gene stacking approach with inducible genes.

This study evaluated Cre-*lox* mediated site-specific integration of a set of 5 genes consisting of 3 constitutively-overexpressed and 2 inducible genes in the crop model rice by analyzing transformation efficiency and expression stability over three generations. High transformation efficiency of the multigene construct harboring repeat sequences, strong expression of the constitutively-expressed genes, and proper regulation of the inducible genes demonstrates that Cre-*lox* mediated integration into the engineered target sites is a practical approach for gene stacking and trait engineering into the crop genomes.

## Results

### Molecular strategy

This study used the Cre-*lox* mediated site-specific integration approach based on the use of mutant *lox* sites, *lox75* and *lox76*, to stabilize the integration locus as described earlier (Albert et al., 1995; Srivastava and Ow, 2002). Specifically, the rice target line, T5, is retransformed by the donor vector pNS64 to obtain the multigene site-specific integration locus. *T5* target locus consists of a *lox76* placed between the maize ubiquitin-1 promoter (*ZmUbi1*) and *cre* coding sequence (**Fig. 1a**), and pNS64 contains a *loxP* and *lox75* -flanked gene construct consisting of a promoter-less selectable marker (*NPT II*) and 4 genes-of-interest (*GFP, GUS, DREB1a, pporRFP*), each expressed by their dedicated promoters (**Fig. 1b**). Upon entry into T5 cells, pNS64 undergoes Cre-*lox* recombination leading to site-specific integration of the gene construct (without vector backbone) into the target site (**Fig. 1c**). The resulting site-specific integration (SSI) is selectable due to the placement of *NPT II* downstream of strong promoter (*ZmUbi1*), and characterized by distinct left and right junctions.

**Figure 1:**
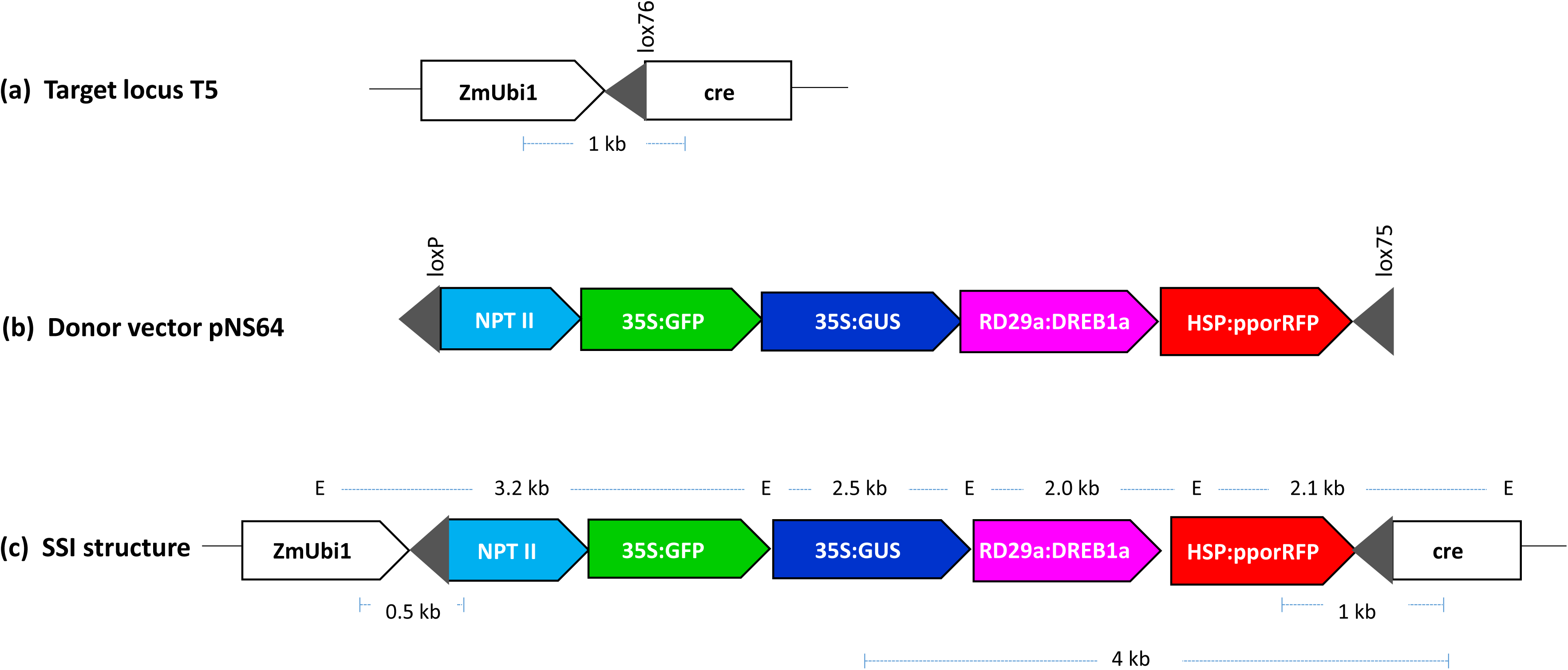
Molecular approach for site-specific integration (SSI) of a multigene stacking. **(a)** *T5* locus in rice cv. Taipei-309 consisting of a single-copy of T-DNA encoding Cre activity and the target *lox76* site (black triangle). **(b)** Donor vector, pNS64, in pBluescript SK backbone (not shown) containing promoterless *NPT II* gene and four expression units (*GFP, GUS, AtDREB1A*, and *pporRFP*) between *loxP* and *lox75* (black triangles). The *loxP x lox75* recombination circularize the gene construct, which subsequently integrates into *T5* locus to generate the site-specific integration (SSI) structure. The *NPT II* gene captures the maize ubiquitin-1 promoter (*ZmUbi1*) at *T5* locus to make the event selectable on geneticin^TM^, and SSI locus expresses four genes, two constitutive (*GFP* and *GUS*) and two inducible (*AtDREB1A* and *pporRFP*) genes. **(c)** Structure of the predicted site-specific integration (SSI) locus that expresses a stack of four genes (*NPT II, GFP, GUS, AtDREB1A*, and *pporRFP*). The primers sites and the expected PCR products are indicated below the structure, while *Eco*RI sites (E) and the fragment sizes are shown above the structure. *35S*: Cauliflower Mosaic Virus *35S* promoter, *NPT II*: neomycin phosphotransferase II, *GFP*: green fluorescent protein, *GUS*: β-Glucuronidase, *AtRD29a*: *Arabidopsis thaliana RD29a* promoter, *AtDREB1A*: *Arabidopsis thaliana* dehydration responsive element 1A, *HSP17.5E*: soybean heat-shock 17.5E gene promoter, and *pporRFP*: sea coral *Porites porites* red fluorescent protein. Each gene carries a nopaline synthase (*nos 3’*) transcription terminator (not shown).

### Characterization of the transgenic lines

Using gene gun mediated transformation of T5 line (cv. Taipei-309) with pNS64, 32 T0 plants were obtained (**Table S1**). Four of the T0 plants were removed from the analysis due to their weak stature at the young vegetative stage, and remaining 28 were analyzed by PCR. Twenty-seven of these lines were found to contain the SSI junctions indicated by 0.5 kb and 1 kb left and the right junction bands, respectively (**Fig. S1a**). Five of these lines contained biallelic integrations as indicated by target site PCR and genetic segregation among T1 progeny (**Fig. S1a; Table S1**). Southern blot hybridization of *Eco*RI-digested genomic DNA showed the presence of predictable 3.2 kb left junction and 2.1 kb right junction, upon hybridization with *GFP* and *pporRFP* probes, respectively (**Fig. S2a-b**), and hybridization with *GUS* and *AtDREB1a* probes, in a subset of 22 lines, showed the predictable 2.5 kb *GUS* or 2.0 kb *AtDREB1a* in all except 4 lines (**Fig. S2c-d**). Thus, full-length SSI copy was found in 18 lines, and the remaining 4 contained truncation of the middle portion of the cassette (**Table S1**). Long PCR (4 kb) at the right junction verified this finding and identified another line (#4) with full-length SSI (**Fig. S1b; Table S1**). The lines lacking *GUS* and *AtDREB1a* genes in the Southern blot failed this PCR, while the remaining showed the predicted 4 kb amplicon, indicating a full-length SSI structure (**Fig. S1b**). In addition, Southern analysis revealed that 13 lines contained only the SSI copy, whereas the remaining contained additional 1 – 3 copies (**Fig. S2; Table S1**). Two clonal lines showing identical hybridization patterns with 4 genes were also identified (**Fig. S2; Table S1**). In summary, the presence of multigene SSI locus was validated in 19 of the 30 SSI lines (removing two clonal line from the initial total of 32) developed from the bombardment of 30 callus plates. Eleven of these lines were selected, based on plant vigor and seed abundance, for gene expression analysis (**Table S1**).

### Expression of the constitutive genes

Three constitutively-expressed genes, *NPT II, GFP*, and *GUS*, in the order of arrangement in the SSI locus, are each controlled by a strong constitutive promoter (**Fig. 1**). While *NPT II* serves as the selectable marker, *GFP* and *GUS* represent the genes-of-interest. Expression of these genes was determined at transcript and protein levels in the T0 lines and their progeny using T5 line as the negative control. Transcript analysis by RT-qPCR, in the T0 plants, showed that most lines abundantly expressed the three genes within 2 to 5-fold range (**Fig. 2a-c**). However, SSI line #12 behaved atypically as it expressed *NPT II* and *GFP* transcripts within the range but showed markedly lower levels of *GUS* transcripts (**Fig. 2c**). Although, gene expression at transcript levels in SSI lines has not been evaluated in previous reports, >10-fold lower GUS transcript in line #12 was somewhat surprising. Transcript abundance analysis in the T1 progeny was performed in 10-day old seedlings grown in the germination media. This analysis showed a greater variation in the transcript levels of the three genes (**Fig. 2d-f**); however, all T1 plants abundantly expressed the three genes. Notably, T1 progeny of line #12 showed a similar pattern as found in the parental line, consisting of higher levels or *NPT II* and *GFP* transcripts but markedly lower levels of *GUS* transcripts (**Fig. 2f**).

**Figure 2:**
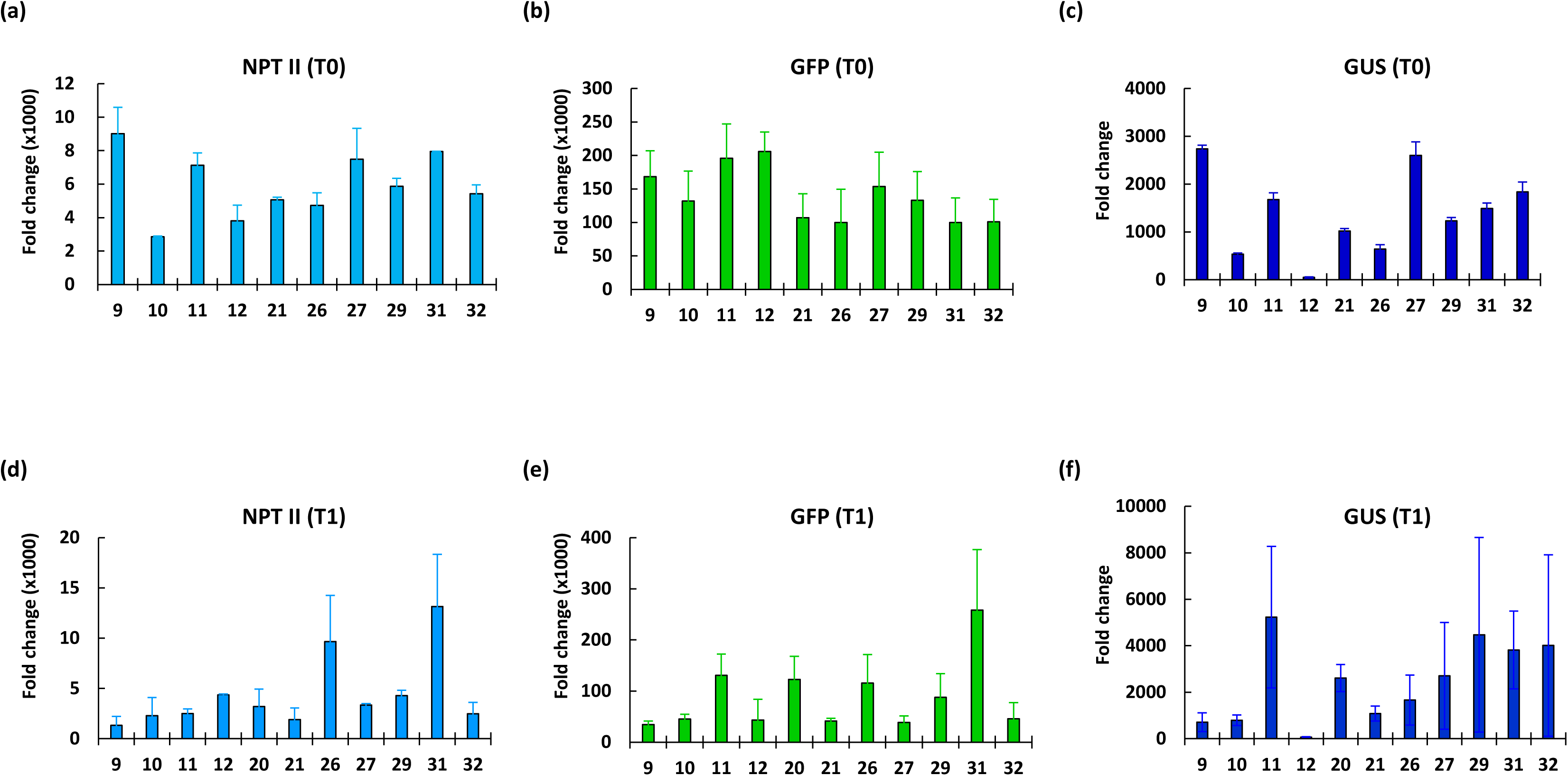
Transcript abundance analysis of constitutively expressed genes by real time quantitative PCR (RT-qPCR). Relative expression of *NPT II, GFP*, and *GUS* genes in the T0 plants (**a – c**) and the T1 progeny seedlings (**d – e**) of site-specific integration (SSI) lines in comparison to the T5 negative control. The SSI line numbers are given on x-axis. The values are the average of 2 biological replicates and 2 technical replicates of each. Standard errors are indicated as the error bars.

Steady state mRNA levels can only partially predict protein abundance as post-transcriptional regulation and cell perturbation among other factors influence the correlation of transcript and protein abundance (Vogel and Marcotte, 2012). Therefore, gene expression analysis at the protein level is important for interpreting the functional effects. In this study, NPT II ELISA, GFP fluorescence, and GUS activity were measured to understand gene expression from the stacked locus. The NPT II activity in T0 plant was expected be strong owing to the selection during tissue culture; therefore, NPT II ELISA was done only in T1 progeny. We found a 4-fold variation in GFP fluorescence and 2.8-fold variation in GUS activity among SSI lines (**Fig. 3a-b**), indicating a good correlation of these protein activities with their transcript levels. However, line #12, to our surprise, showed a comparable GUS activity (**Fig. 3b**), even though it was found to express markedly lower levels of the GUS transcripts (**Fig. 2c**). The reason behind this discrepancy is not clear but it could be an artifact of the GUS assay. If optimum amount of substrate is not used in the reaction, the enzyme rate may not be directly proportional to the enzyme concentration (Robinson, 2015). To further check GUS activity in line #12, histochemical staining of leaf cuttings was done according to Jefferson et al. (1987). A lower staining intensity in line #12 compared to other lines such as #10 (**Fig. 3c**), suggested that #12 contains a lower GUS activity, and the transcript abundance in this line is a good indicator of its GUS levels. Among T1 progeny of the SSI lines, only <3-fold variation for NPT II and GFP abundance, and ∼5-fold variation for GUS activity was observed (**Fig. 3d-f**). Finally, a subset of 4 SSI lines was selected for T2 progeny analysis. T2 seedlings of these lines were found to abundantly expressed GFP fluorescence and GUS activities at more or less similar levels (**Fig. 3g-h**), indicating that stacked genes at the SSI locus are stably transmitted to the progeny. The stability of the genes was also reflected by the gene dosage effect as the biallelic lines generally expressed GFP and GUS at higher levels compared to the monoallelic lines, although, a significant difference was only found for GFP (**Fig. 3i**). In summary, stable expression of the three stacked genes regulated by strong promoters at the SSI locus was found in all lines tested in this study. Only one SSI line (#12) showed sub-optimal expression of one of the three genes (GUS), while all other showed comparable expression that were transmitted to the subsequent generations.

**Figure 3:**
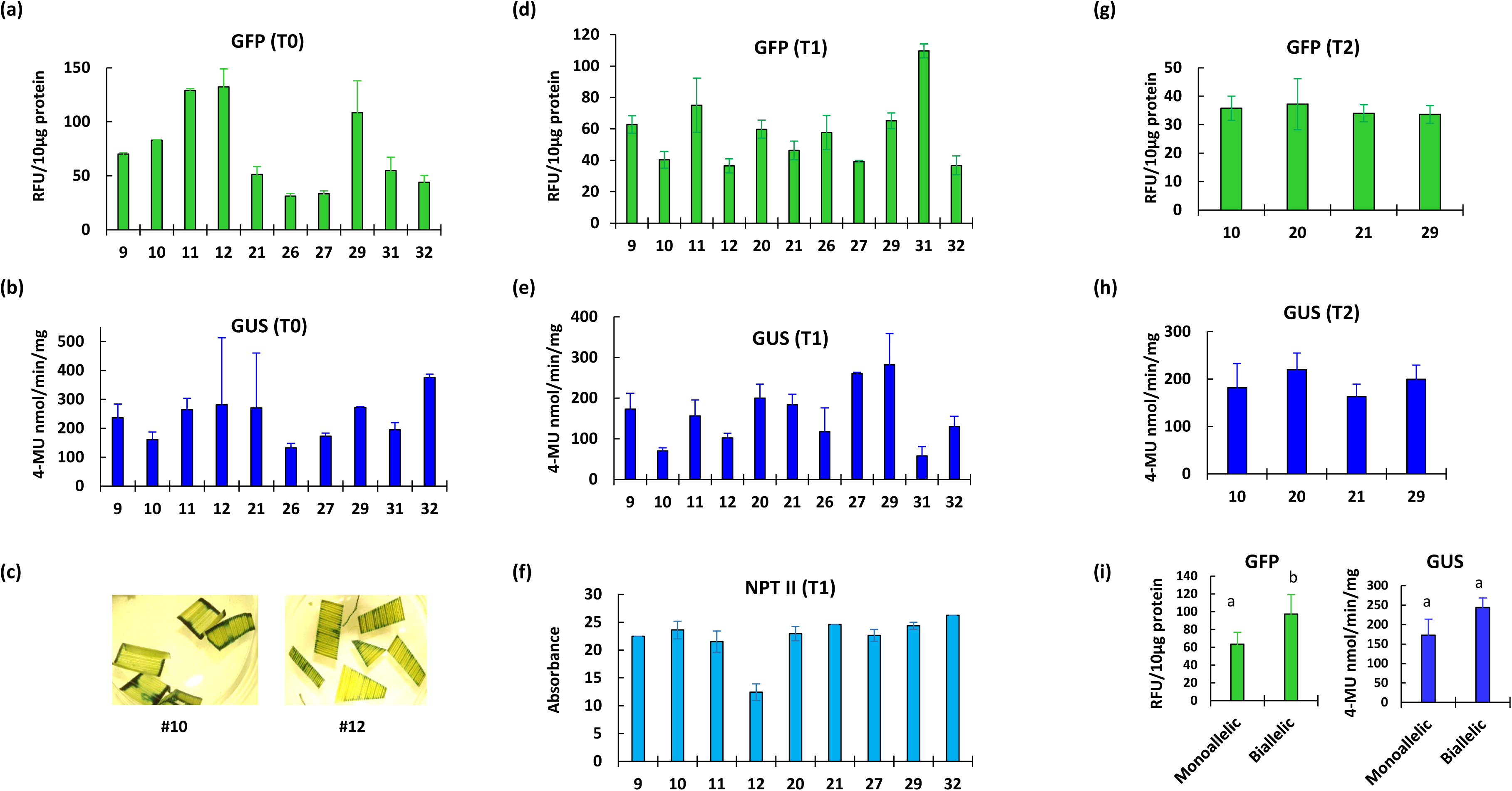
Protein abundance analysis of constitutively-expressed genes in the site specific integration (SSI) lines. (**a – b**) Quantitative GFP fluorescence and GUS activity in T0 plants. (**c**) comparative histochemical GUS staining of leaf cuttings from SSI lines #10 and #12. (**d – f**) GFP fluorescence, GUS activity, and NPT II ELISA in T1 progeny seedlings. (**g – h**) quantitative GFP fluorescence and GUS activity in T2 progeny seedlings. (**i**) average GFP and GUS activities in monoallelic and biallelic T0 SSI lines (described in Table S1). The SSI line numbers are given on x-axis. Statistical differences, shown by the alphabets, were determined by student *t*-test at *p*=0.05. Standard errors are indicated as the error bars.

### Expression of inducible genes

The SSI locus contained two inducible genes, a transcription factor, *AtDREB1a*, and a fluorescent protein, *pporRFP*, driven by *RD29a* and *HSP17.5E* promoters, respectively. These genes are expected to be ‘off’ at room temperature but abundantly induced by specific treatments, *i.e*., cold-shock for *RD29a* and heat-shock for *HSP17.5E*. The expression pattern of these genes was studied by RT-qPCR in T0 plants and their progeny, at room temperature and upon treatment, using T5 as the negative control. Expression analysis of *AtDREB1a* in T0, T1, and T2 plants showed abundant transcripts upon cold-shock treatment (**Fig. 4a – c**). Similarly, *ppor*RFP analysis in T0, T1, and T2 plants showed abundant transcripts in heat-shock treated samples (**Fig. 4d – f**). These experiments verified that the two inducible genes work properly and respond to the treatment. Notably, both genes expressed at a basal level at the room temperature, indicating proper regulation of the genes in SSI lines, and its faithful transmission to the progeny. The fold-induction was highly variable between lines, but most lines showed strong induction upon treatment. Finally, specificity of the genes was determined by checking their expression upon non-specific perturbation, *i.e*., induction of *RD29a:AtDREB1a* by heat-shock and *HSP17.5E:pporRFP* by cold-shock treatment. As expected, heat-shock treatment had little effect on *AtDREB1a* and cold-shock treatment did not induce *pporRFP* (**Fig. S3a-b**), showing that stacking these two genes closely together did not break their promoter-specificity in the SSI lines.

**Figure 4:**
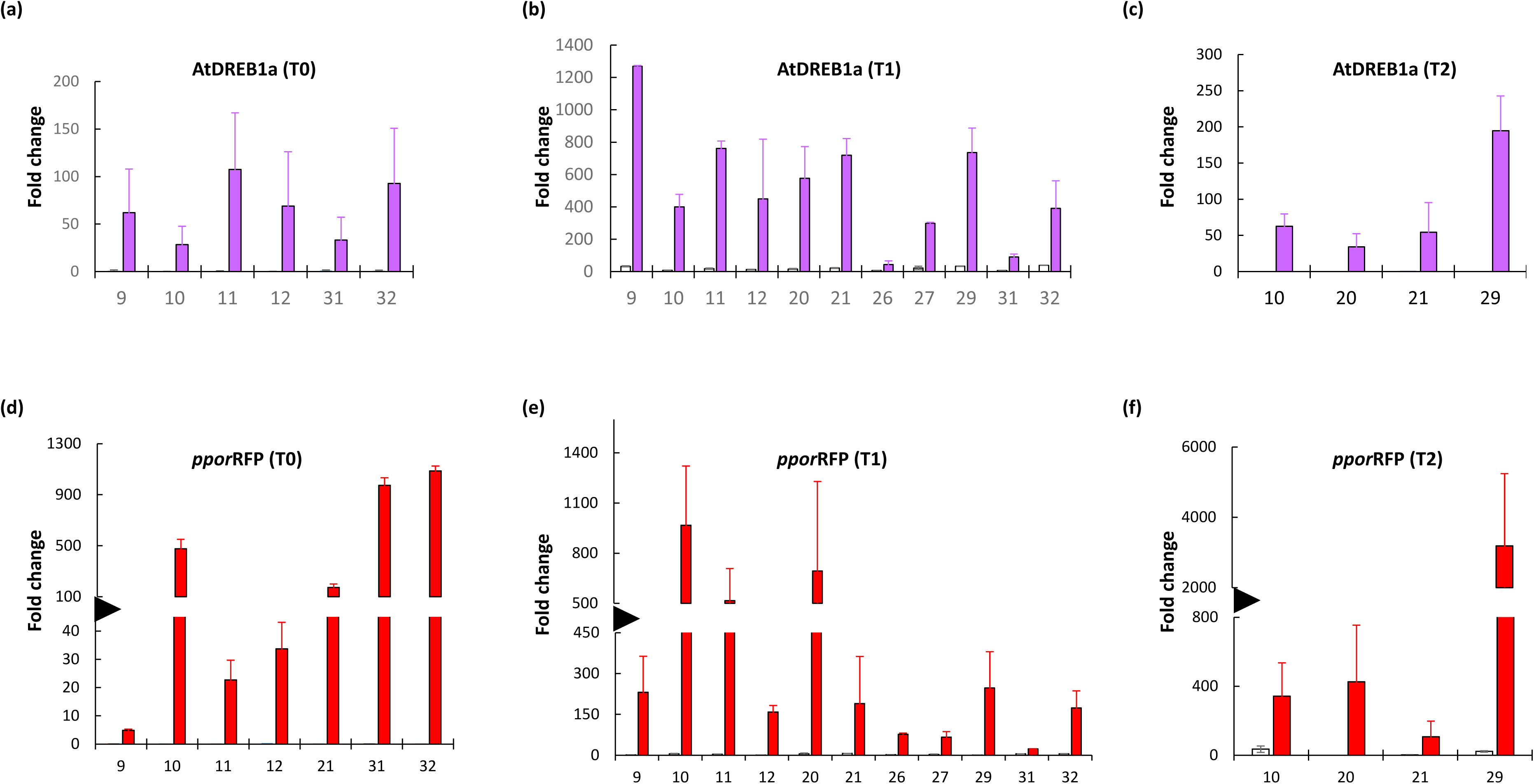
Expression analysis of the inducible genes by real time quantitative PCR (RT-qPCR) in the site-specific integration (SSI) lines relative to T5 negative control. (**a – c**) *AtDREB1a* expression analysis at room temperature (white bars) or upon cold-induction (20 hours in ice pack, magenta bars) in T0, T1, and T2 plants. **(d – f)** *pporRFP* expression at room temperature (white bars) or upon heat-induction (42°C for 3 hours; red bars) in T0, T1, and T2 plants. The SSI line numbers are given on x-axis. The values are the average of 2 biological replicates with standard error indicated as the error bars.

Downstream targets of *AtDREB1a* transcription factor during cold stress are not clearly known in rice (Oh et al., 2005); therefore, it’s downstream effects in SSI lines could not be analyzed. The functional effects of AtDREB1a expression would require phenotypic analysis in stress conditions. Therefore, protein abundance analysis was performed only on *pporRFP*. Confocal imaging of root or leaf samples was done on the T1 seedlings of three SSI lines (#9, #10, #12) using GFP as the internal control. *ppor*RFP is a complex dsRed type RFP that undergoes homo-dimerization to become functional and emit fluorescence (Saccheetti et al., 2002). In a constitutive expression system, dimerization would occur continuously. However, in the inducible expression system, it is important to determine the lag time for *ppor*RFP maturation. For this, SSI #9 seedlings were heat-treated and their roots were imaged at 3 different time intervals. Optimum fluorescence was observed 72 hours after heat-treatment (**Fig. S4)**. This is in agreement with the published report on temporal decoupling of dsRed RFP mRNA and protein expression. In Medicago, dsRed RFP mRNA was found to peak at 2 – 3 days after transformation, while fluorescence peak occurred 3 – 5 days later (Jansing and Buyel 2019). Accordingly, seedlings of SSI lines #9, #10, and #12 were subjected to 3 h of heat-shock treatment at 42°C followed by fluorescence imaging 72 h later in confocal microscope. As expected, each SSI line abundantly expressed GFP regardless of the treatment. Whereas, *ppor*RFP showed an inducible pattern, characterized by undetectable red fluorescence at room temperature (RT) and enhanced fluorescence upon heat-shock (HS) in the root or leaf tissues (**Fig. 5**). In summary, the two co-localized inducible genes that are regulated by specific stress treatments, expressed properly from the stacked SSI locus without apparent interference of the neighboring strong constitutive genes or the inducible genes.

**Figure 5:**
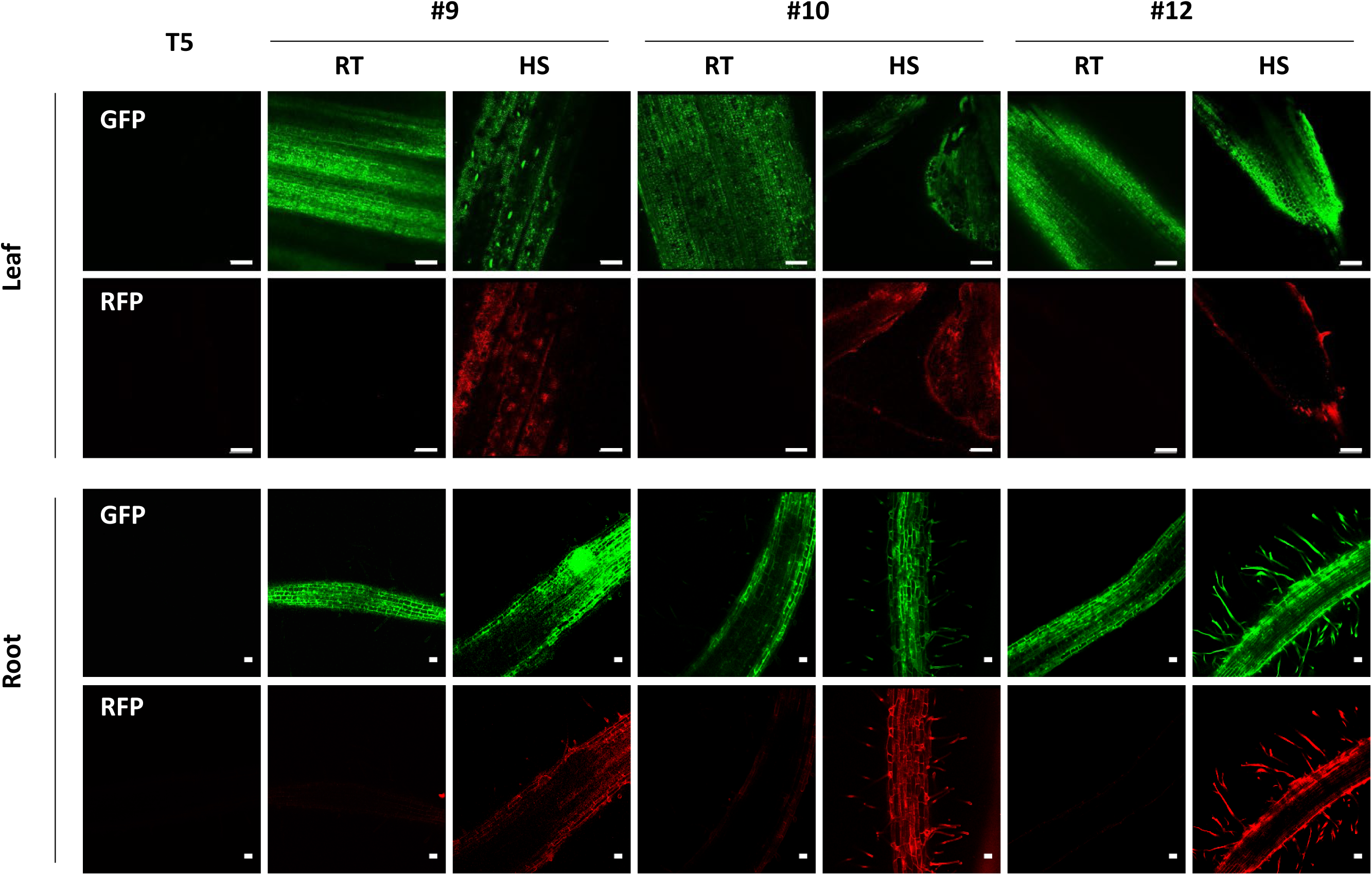
Confocal imaging of GFP (top) and *ppor*RFP (bottom) in the roots and leaves of the 10 days old T1 seedlings of site-specific integration lines #9, #10, and #12. All images were taken 72 hours post heat-shock treatment at 20x magnification. T5: Target line (negative control); RT: Room Temperature; HS: Heat-shock. Scale bar in leaves: 100 µm and in roots: 50 µm.

## Discussion

Multigene transformation has overcome the technical barriers of high load gene transfer but the challenges related to co-integration and co-expression remain to be fully addressed. Both *Agrobacterium* and gene gun are competent at multigene transfers; however, complexity of DNA integration into the plant cell leads to multiple random outcomes consisting of complex integrations into unique genomic sites (Saika et al., 2017; Somers and Makarevitch, 2004). Co-bombardment of gene vectors or multi-genic T-DNA have been used for metabolic engineering in plants (Naqvi et al., 2009; Ruiz-Lopez et al., 2015), however, low rate of co-expression from multigene assemblies poses a major bottleneck (Ghareeb et al., 2016; Schmidt et al, 2008). Recombinase-mediated site-specific integration overcomes the complexity of DNA integration and simplifies the structure, which in turn removes gene silencing triggers and creates a favorable environment for stable gene expression (Akbudak et al., 2010; Chawla et al., 2006; Day et al., 2000; Li et al., 2010; Nanto et al., 2009). In this study, we evaluated the recombinase approach for multigene transformation, specifically, asking how efficiently a multigene construct integrates into the targeted site and whether the integrated genes express faithfully in the plants. Using an available target line, we developed site-specific integration of a 5-gene cassette through Cre-*lox* recombination in rice. The molecular strategy of site-specific integration has been described earlier (Srivastava and Ow, 2002). Using gene gun mediated particle bombardment, 30 transgenic events were obtained from 30 shots, 19 of which contained precise multigene integration characterized as full-length integration of the cassette into the target site. This efficiency of co-transformation is exceptionally higher than generally found in conventional methods. As observed in previous studies (Albert et al., 1995; Srivastava et al., 2004), the majority of these lines (13 out of 19) contained only the SSI copy, and random integrations were low or undetectable. Most importantly, all tested lines expressed each of the 5 genes, affording stable expression at 100% rate for the multigene assembly in this study. The strong constitutive genes showed abundant expression levels and the inducible genes functioned according to their promoter specificity. Notably, the two inducible genes showed no apparent interference of the neighboring genes. Additionally, the multigene cassette was faithfully transmitted to the progeny as shown by gene expression analysis in T1 and T2 progeny. Interestingly, the repeated use of *nos* terminator in the construct did not seem to affect the structure stability. However, for product development, this feature should be avoided as repeat sequence in multigene assemblies could induce homology-directed truncations. In conclusion, recombinase-mediated site-specific integration approach proved to be highly efficient in developing site-specific integration lines harboring precise integration of the 5-gene cassette, and the efficacy of the approach was further demonstrated by proper expression of all genes and their inheritance by the progeny. The recombinase mediated multigene transformation approach can also be practiced with the ‘cassette exchange’ strategy as described earlier (Li et al., 2010). In addition, the strategy can easily be modified to incorporate marker-removal feature by employing another recombinase system (Nandy et al., 2012), that have been found to function in plant cells (Chen and Ow, 2017; Cody et al., 2020).

## Acknowledgement

We thank Soumen Nandy for pNS64 construction, Shan Zhao for rice transformations, Betty Martin for confocal imaging, and Jessica Kivett for the greenhouse support. This project was supported in part by USDA-NIFA grant 2017-38821-26412 and NSF-EPSCoR grant 1826836.

## Experimental Procedures

### Vector construction and transformation

The multigene vector, pNS64 (Fig. 1b), was developed through the standard restriction digestion and ligation method. The individual gene cassettes from pUC vectors were ligated one by one into pAA12 backbone that contains a promoterless neomycin phosphotransferase II (*NPT II*) gene between *loxP* and *lox75* (Akbudak and Srivastava, 2017). Two gene cassettes, 35S:GFP and 35S:GUS were already available that contained CaMV 35S promoter driving GFP or GUS gene. To build RD29a:AtDREB1a cassette, *Arabidopsis thaliana* dehydration responsive element B1A (*AtDREB1A)* and *Arabidopsis* cold inducible *RD29a* promoter sequences were generated by PCR. For HSP17.5E:pporRFP cassette, the promoter fragment of *Gmhsp17.5E* was obtained by PCR on soybean DNA and *ppor*RFP from pANIC6A that contains coral *Porites porites RFP* (Alieva et al. 2008; Mann et al., 2012). Primers used for cloning are given in **Table S2**. All gene constructs contain nopaline synthase transcription termination sequence (*nos 3’*).

The rice line T5 (Taipei 309) that contains a Cre-*lox* target site was used in the present study to develop site-specific integration lines. The development of T5 line is described by Srivastava and Ow (2002). The pNS64 vector was delivered by gene gun (PDS 1000, Bio-Rad Inc.) into the embryogenic callus derived from T5 mature seeds. The bombarded callus was selected on 100 mg/l geneticin™to isolate the site-specific integration lines. The tissue culture media for callus induction and plant regeneration were used according to Nishimura et al. (2006).

### Molecular analysis

The polymerase chain reaction (PCR) was performed on genomic DNA using Emerald Amp MAX PCR Master Mix (Takara Bio, CA, USA) using the primers given in **Table S2**. Southern blot analysis was performed on genomic DNA digested with *Eco*R1 using ^32^P-labeled DNA probes of GFP, RFP, GUS, and AtDREB1A. Gene expression analysis by reverse transcriptase (RT)-quantitative (q) PCR was performed on total RNA isolated using Trizol reagent (Invitrogen, Inc.) according to manufacturer’s protocol, and quantified on Nano-drop 2000 (Thermo-Fisher Inc.). Two microgram of total RNA, treated with RQ1-RNAse free DNase (Promega Inc.), was used for the cDNA synthesis using PrimeScript RT reagent kit (Takara Bio, CA, USA), and used for qPCR using TB green Premix Ex Taq II (Takara Bio, CA, USA) on Bio-Rad CFX 96 C1000. The product specificity was verified by the melt curve analysis and the Ct values were normalized against 7Ubiquitin. The relative expression was calculated against T5 negative control and the untreated controls using delta-delta Ct method (Livak and Schmittgen, 2001). Each line contained two to three biological replicates with two technical replications of each. For inducible gene expression analysis, excised leaf blades from the greenhouse plants wrapped in aluminum foil or seedlings germinated on MS/2 media in petri dishes, were placed on ice pack for 20 h for cold-shock treatment or 42°C incubator for 3 h for heat-shock treatment. The room temperature controls were handled in the same way except they were kept at the ambient room temperature.

### Protein Analysis

NPT II enzyme linked immunosorbent assay (ELISA) were conducted using a commercial kit (Agdia Inc. USA) according to the manufacturer’s instructions. Fifty milligrams of fresh leaf from 1-month old plants was ground in the protein extraction buffer provided in the kit, and centrifuged to collect the crude protein extract. The NPT II protein provided in the kit and the T5 crude protein extract were used as a positive and negative controls, respectively. ELISA plates were read at A650 in the Synergy Biotek Cytation 3. The ratio of the absorbance of samples to T5 negative control was used as the measure of NPT II expression.

GFP fluorescence was observed in the young tissue under the Leica 56D stereoscope fitted with the 440 – 460 nm excitation and 500 – 560 nm (band pass) emission filters (Night Sea, Lexington, MA). For the quantitative estimation, fresh tissue was ground in 10 mM Tris–EDTA, pH 8.0, at 4°C and centrifuged at 13,000 rpm for 20 min to collect the supernatant. A strongly expressing rice GFP line, C30-1, described earlier (Pathak et al., 2019), was used as a reference. Ten microgram of protein was used for GFP fluorescence measurement in Versa-fluorimeter (Bio-Rad Inc) equipped with 490 ± 5 nm excitation filter and 510 ± 5 nm emission filter. All lines were measured against C30-1 positive control and the T5 negative control. A unit of GFP was defined as relative fluorescence units per 10 µg of total protein (RFU/10 µg).

Histochemical GUS staining was done by submerging leaf cuttings in the GUS staining solution containing 1 mM X-Gluc (Gold Biotechnologies, St. Louis, MO, USA) according to Jefferson et al. (1987). For quantitative measurements, 10 µg of total protein extracted in 50 mM phosphate buffer, pH 7, was used for GUS assay using 4-methylumbelliferyl b-D-glucuronide (4-MUG) as the substrate, and the kinetics of the reaction measured in spectrophotometer by appearance of a colored product 4-methylumbelliferone (4-MU) according to Jefferson et al., (1987). A unit of GUS activity was defined as nmol 4-MU produced per minute from 1 mg of the protein (nmol/min/mg). For each of the above assays, 2 – 3 biological replicates with the two technical replicates were included, and all protein estimations were done using Bradford reagent (VWR Inc.).

### Confocal Imaging

The *ppor*RFP detection was performed using confocal imaging in the 7 – 10 days old seedlings. The seedlings were heat-shocked as described in the molecular analysis section, and imaging was done at 24, 48 and 72 hours post heat-shock treatment. The images were captured using a Leica TCS SP5 (Buffalo Grove, IL. USA) confocal microscope by the bandwidth adjustment for the fluorescence detection. For roots imaging, the samples were excited using 514 Argon and 594 HeNe laser channels and emission was collected at 542 – 582 mm for GFP, and at 610 – 710 nm for *ppor*RFP. For leaf imaging, samples were excited at 514 Argon laser channel, and emission was collected at 590 – 610 nm for blocking the chlorophyll auto-fluorescence. The leaf images were captured through sequential scan to prevent the bleed-through between chlorophyll auto-fluorescence and fluorescent protein(s). A GFP positive line, C30-1, described earlier (Pathak et al., 2019), and the parental T5 seedlings were used as controls. For all samples, first the gain, zoom and offset were adjusted for T5 negative control, and then all images were captured using the same parameters at 20x magnification.

---

**Supplementary Figure 1**: PCR verification of 28 T0 site-specific integration (SSI) lines. (**a**) presence of left SSI junction, right SSI junction, and the target site. (**b**) Long PCR at right junction to verify full-length integration. Primer positions in the SSI and target sites, and the fragment sizes are shown in Figure. 1. T5, target line as negative control; NTC, no template control.

**Supplementary Figure 2:** Southern hybridization of *Eco*R1-digested genomic DNA of T0 site-specific integration (SSI) lines using (**a**) *GFP*, (**b**) *pporRFP*, (**c**) *GUS* and, (**d**) *AtDREB1A* probes. Size markers are indicated. *Eco*R1 map and fragment sizes of the SSI structure is shown in Fig. 1.

**Supplementary Figure 3**: Non-specific perturbation analysis of the inducible genes by real time quantitative PCR (RT-qPCR) on the T1 progeny seedlings of 3 site-specific integration (SSI) lines relative to the T5 negative control. (**a**) *AtDREB1a* analysis, and (**b**) *pporRFP* analysis in heat-shock (HS) and cold-shock (CS) treated samples. Standard errors are indicated as the error bars.

**Supplementary Figure 4**: Time course confocal imaging of *ppor*RFP in 10 days old T1 seedlings of SSI line #9. Images are captured 24, 48 and 72 hours post heat-shock treatment at 20x magnification using the same parameters. GFP imaging is included as an internal control. T5, negative control; RT, room temperature.

