## Supplementary figures and images for "Recombinase-mediated integration of a multigene cassette in rice leads to stable expression and inheritance of the stacked locus"

### Fig. S1 - S4

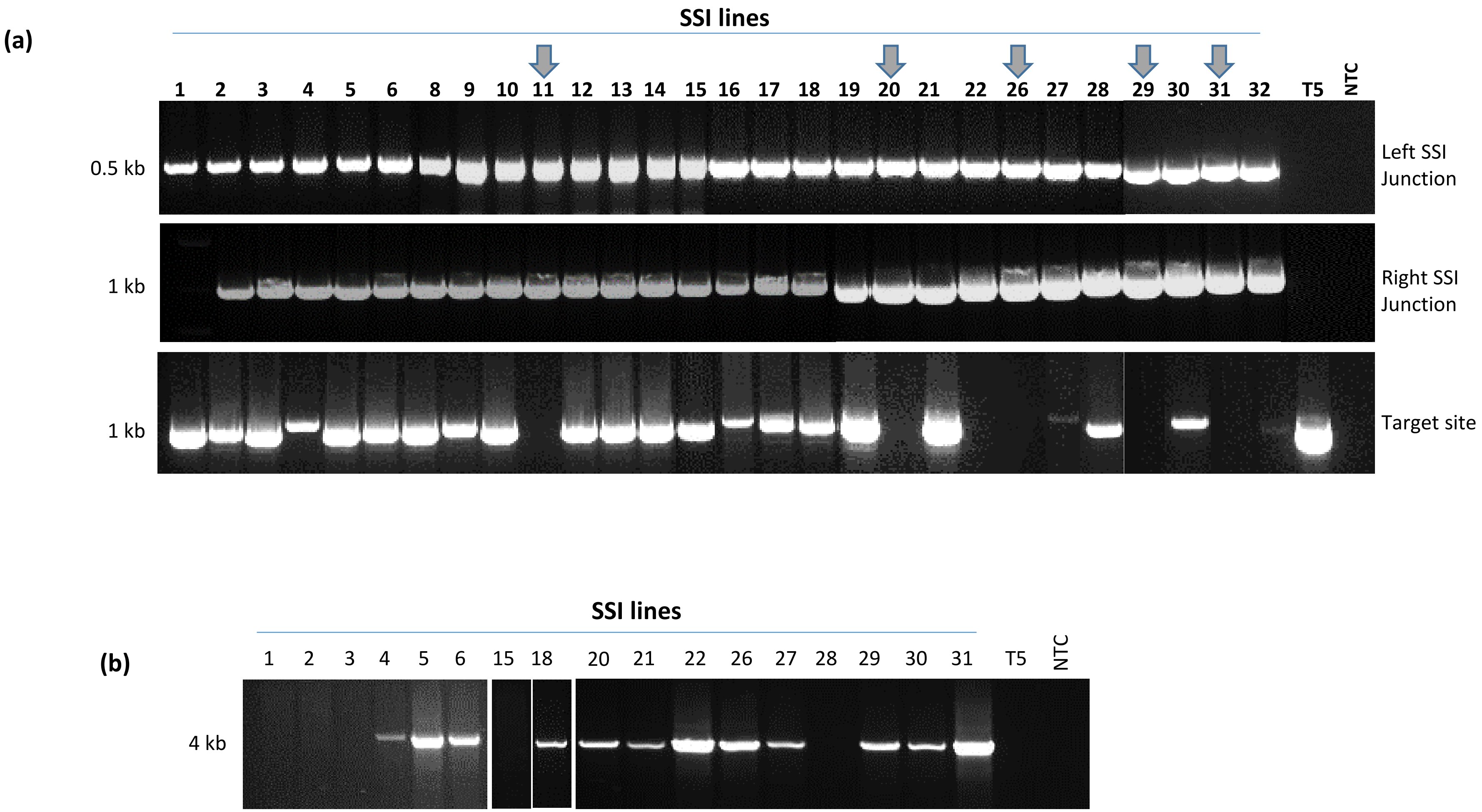

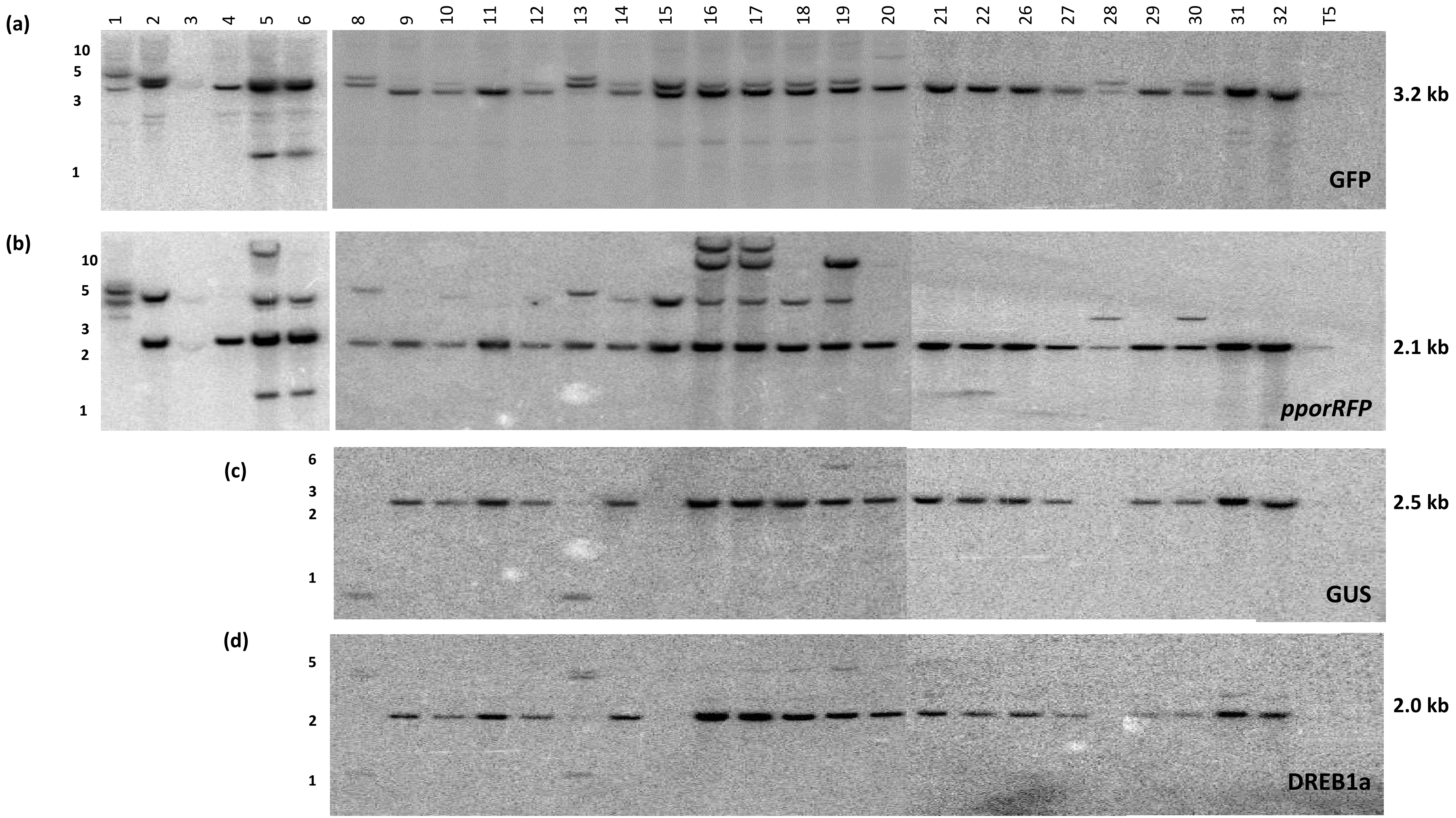

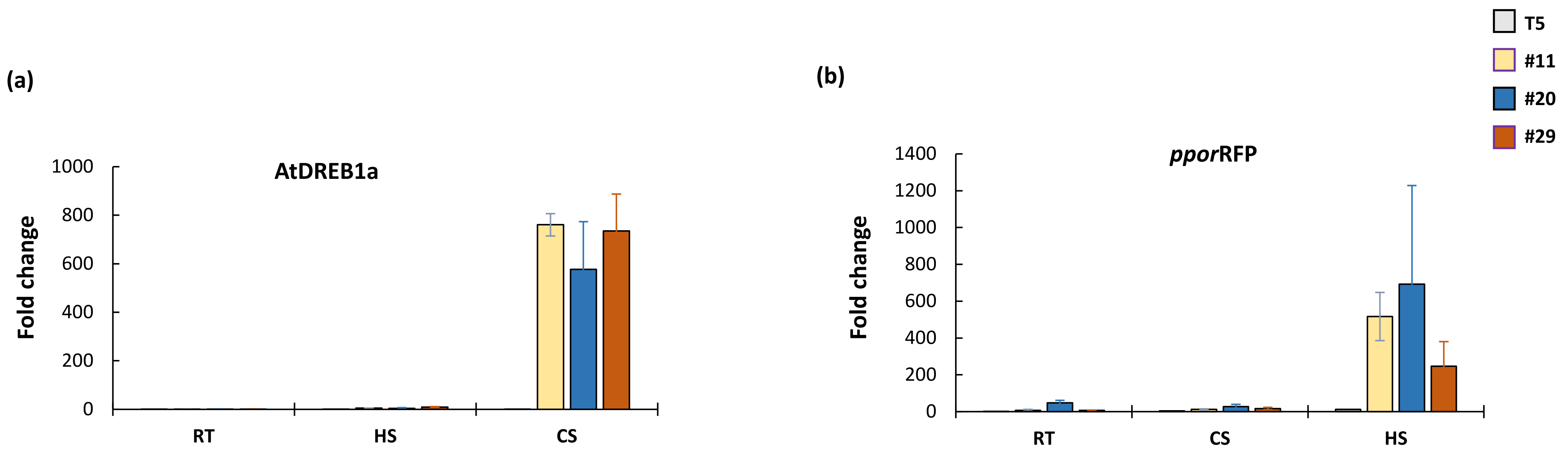

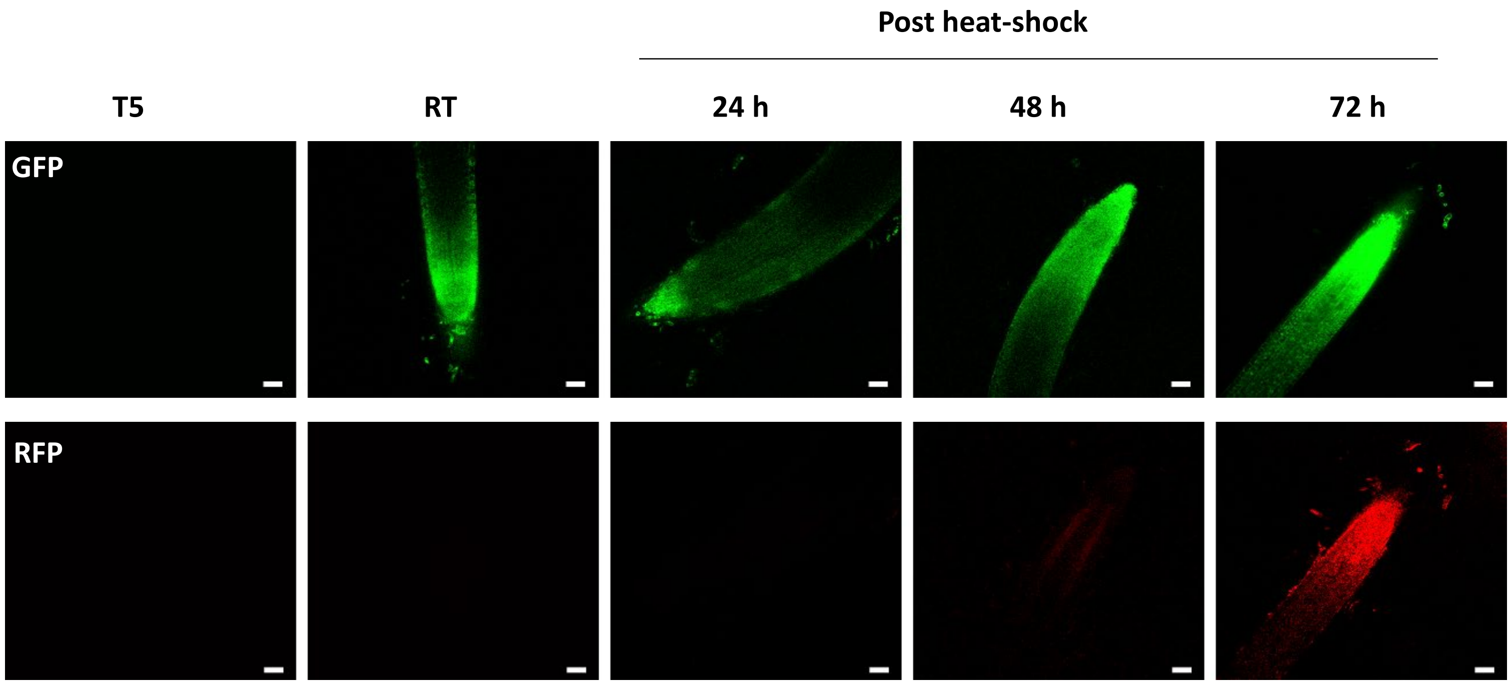


Fig. S4
