## Supplementary material for "Recombinase-mediated integration of a multigene cassette in rice leads to stable expression and inheritance of the stacked locus": Table S1

| **Lines** | **Site-specific integration^1^** | **Full-length integration** | **Random integrations^2^** | **T1 Segregation** | | **Conclusion^3^** | **Selected for analysis** |
| --- | --- | --- | --- | --- | --- | --- | --- |
|  |  |  |  | **GFP+** | **GFP-** |  |  |
| 1 | - | - | - | - | - | Non-SSI |  |
| 2 | ✓ | - | ✓ | - | - | Truncated |  |
| 3 | ✓ | - | ✓ | - | - | Truncated |  |
| 4 | ✓ | ✓ | - | - | - | Weak plant |  |
| 5 | ✓ | ✓ | ✓ | 8 | 2 | Monoallelic |  |
| 6 | ✓ | ✓ | ✓ | - | - | Clonal to 5 |  |
| 7 | ND | ND | ND | - | - | - |  |
| 8 | ✓ | - | ✓ | - | - | Truncated |  |
| 9 | ✓ | ✓ | - | 44 | 13 | Monoallelic | ✓ |
| 10 | ✓ | ✓ | - | 38 | 5 | Monoallelic | ✓ |
| 11 | ✓ | ✓ | - | 47 | 0 | Biallelic | ✓ |
| 12 | ✓ | ✓ | - | 26 | 14 | Monoallelic | ✓ |
| 13 | ✓ | - | ✓ | - | - | Truncated |  |
| 14 | ✓ | ✓ | ✓ | 16 | 6 | Monoallelic |  |
| 15 | ✓ | ✓ | ✓ | - | - | Truncated |  |
| 16 | ✓ | ✓ | ✓ | 25 | 4 | Monoallelic |  |
| 17 | ✓ | ✓ | ✓ | - | - | Clonal to 16 |  |
| 18 | ✓ | ✓ | ✓ | - | - | Monoallelic |  |
| 19 | ✓ | ✓ | ✓ | 13 | 2 | Monoallelic |  |
| 20 | ✓ | ✓ | - | 23 | 0 | Biallelic | ✓ |
| 21 | ✓ | ✓ | - | 23 | 5 | Monoallelic | ✓ |
| 22 | ✓ | ✓ | - | - | - | Biallelic/sterile plant |  |
| 23 | ND | ND | ND | - | - | Weak plant |  |
| 24 | ND | ND | ND | - | - | Weak plant |  |
| 25 | ND | ND | ND | - | - | Weak plant |  |
| 26 | ✓ | ✓ | - | 17 | 4 | Monoallelic | ✓ |
| 27 | ✓ | ✓ | - | 14 | 7 | Monoallelic | ✓ |
| 28 | ✓ | - | - | - | - | Truncated |  |
| 29 | ✓ | ✓ | - | 19 | 0 | Biallelic | ✓ |
| 30 | ✓ | ✓ | ✓ | - | - | Monoallelic |  |
| 31 | ✓ | ✓ | - | 18 | 0 | Biallelic | ✓ |
| 32 | ✓ | ✓ | - | 19 | 6 | Monoallelic | ✓ |

**Table S1: Characterization of T0 lines**

^1^Determined by PCR and Southern blot analysis (Fig. S1, S2); ^2^Presence of extra-SSI bands in Southern hybridization (Fig. S2); ^3^Characterization of T0 plants based on plant vigor, molecular analysis, and genetic segregation; ND, not determined.
