## Supplementary material for "Recombinase-mediated integration of a multigene cassette in rice leads to stable expression and inheritance of the stacked locus": Table S2

**Table S2: Primers used in this study**

| **Primer Name** | **Sequence (*5’ - 3’*)** | **Application** |
| --- | --- | --- |
| Ubi1960 | GCTCACCCTGTTGTTTGGTG | Genotyping of site-specific integration lines |
| KanR | CTCGATGCGATGTTTCGCTT |  |
| pporRFP-F | CGGGATCCATGGCTCTTTCAAAGC |  |
| Cre-R2333 | ATTGCTGTCACTTGGTCGTG |  |
| Ubi-F | TCTACTTCTGTTCATGTTTGT |  |
| creATG-F | ACGGTCAGTAAATTGGACAT |  |
| Gus-F982 | ACCTCGCATTACCCTTACGC |  |
| qDREB-F | GGAGACGTTGGTGGAGGCTA | *AtDREB1A* qPCR |
| qDREB-R | CGGACGGAAGCGGCAAAAGCA |  |
| qRFP-F | GGCTCGATGGCGACTCTTTCAT | *pporRFP* qPCR |
| qRFP-R | CACCACACTCATACAGTCTCT |  |
| qGFP-F | GACCACTACCAGCAGAACAC | *GFP* qPCR |
| qGFP-R | CCATGTGATCGCGCTTCT |  |
| qNPT-F | CGTTGGCTACCCGTGATATT | *NPT* qPCR |
| qNPT-R | CTCGTCAAGAAGGCGATAGAAG |  |
| qGUS-F | CGACCTCGCAAGGCATATT | *GUS* qPCR |
| GUS-R2 | TCACCGAAGTTCATGCCAGT |  |
| Q7Ubiq1445-F | TGGTCAGTAATCAGCCAGTTTG | Reference gene for qPCR |
| Q7Ubiq1520-R | CAAATACTTGACGAACAGAGGC |  |
